# CRF neurons of the BNST promote resilience by blunting the internal experience of aversion

**DOI:** 10.1101/2022.10.16.512419

**Authors:** Sherod E Haynes, Helen S Mayberg, Larry J. Young, Ming-Hu Han

## Abstract

The Bed Nucleus of the Stria Terminalis (BNST) has been studied extensively for its coordination of opposing adaptive behaviors. Previously, we uncovered a critical role for Corticotropin-Releasing Factor (CRF)-expressing neurons of the oval nucleus of the BNST (BNSTov^CRF^) in maintaining resilience to social defeat through stress-dependent self-sustaining neuronal activity.^1^ However, as mice develop resilience, it is not well-understood how affect and motivation are altered to achieve adaptive behavior in the face of ongoing threat. Here, we explore how this neuronal population exerts a powerful influence over internal state in various stress contexts to promote adaptive social responding. Using cell-type-selective optogenetics, a suite of behavioral paradigms, and transgenic Crf-ChR2 mice, we show that BNSTov^CRF^ neurons induce resiliency by altering the encoding of psychosocial stress, enhancing the appetitiveness of social interaction, and enhancing tolerability to physical stress. Adaptive responses to stress typically emanate as a response to negative internal states by external stimuli; here, we show that in resilient mice, stressful environments are less aversive than susceptible mice, suggesting a different motivational capacity to endure stress in this group. Thus, we describe a novel role for BNSTov^CRF^ neurons in resisting the emotional effects of cumulative stress by reducing the internal experience of aversion

## Introduction

The oval nucleus of the Bed Nucleus of the Stria Terminals (BNSTov) serves as a key integrator of intero- and exteroceptive information has important implications for playing a causative role in the development of depression that may be distinct from that which is involved in recurrent or sustained disease states. The subjective experience of an external stimulus is influenced by one’s internal state, called alliesthesia. Internal states are governed by the milieu of interoceptive, viscerosensory, motivational, and homeostatic cues and inputs. The BNST is well-positioned to serve as the hub of this integration, and dysregulation of this region may lead to vulnerability or resiliency to developing MDD^2^.

Social contexts require both the assessment of environment (i.e., aggression, motivation of social target, prediction of danger or aggression, presence of sexual receptivity) and integration of internal state (i.e., anxiety/fear states, the necessity for interaction to achieve homeostatic needs such as feeding or sex), and thus provide a readout of deficits in interoception. Corticotrophin-releasing factor (CRF) neurons of the BNST have been implicated in this integrative capacity by way of connections to the anterior insula and basolateral amygdala (for example). However, what is not well understood is how emotional valence is achieved across stress contexts. CRF neurons have been shown to possess biphasic properties, where in some settings the neuronal population is appetitive and others aversive.

In this report, we interrogate the role of BNSTov^CRF^ neurons in altering internal motivation for pro-social behaviors that define resilience. We first parsed the bivalent role this neuronal population plays as instrumenting aversion in low or no stress and appetition in the setting of chronic stress. Next, we describe the role for BNSTov^CRF^ neurons in increasing sociability toward familiar aggression. Lastly, we demonstrate their role in shifting socio-affecive bias in social settings, as well as blunting aversion in both social and non-social contexts. Taken together, we show that resilience is not an inappropriate attribution of appetitive in potentially hazardous settings, rather a blunting of aversion which thereby promotes behavioral flexibility and adaptive engagement for survival.

## Results

### BNSTov^CRF^ neuronal stimulation in low- or no-stress conditions induces social avoidance

Despite our findings that activation of Corticotropin-Releasing Factor (CRF)-expressing neurons contained within the oval nucleus of the Bed Nucleus of the Stria Terminalis (BNSTov) produces pro-resilient behavioral responses, several studies have reported that this activation induces negative emotional states.^3–8^ Previously, we show that BNSTov^CRF^ activation had a pro-social effect only after cumulative stress exposure^1^, suggesting a role for BNSTov^CRF^ neurons in potentially promoting cooperative sociality under stressful conditions. This adaptive function confers an evolutionary advantage.^9–11^ However, it is unknown what function these neurons serve in low or no stress conditions. While there are no published reports on the role of BNSTov^CRF^ neurons in social settings, we predicted that optical stimulation would produce avoidant responses to an otherwise pro-social appetitive experience in line with published work in non-social settings.^12–15^ To test this, transgenic mice expressing channelrhodopsin exclusively in Crf-expressing neurons (ChR2 mice) mice were implanted with fiber optic ferrules and received 5 Hz stimulation during the social interaction test (Fig 1a). Control mice showed no significant difference in time spent in the interaction zone with and without a social target; however, there was a trend toward more significant time spent with the target animal (p=0.08, t-test two-tailed, Fig 1b). In contrast, ChR2 mice had a significant decrease in time spent with a novel conspecific between the no target and target trials, suggesting that acute BNSTov^CRF^ stimulation induced negative valence onto an otherwise rewarding social context (Fig 1b).

**Figure 1.**
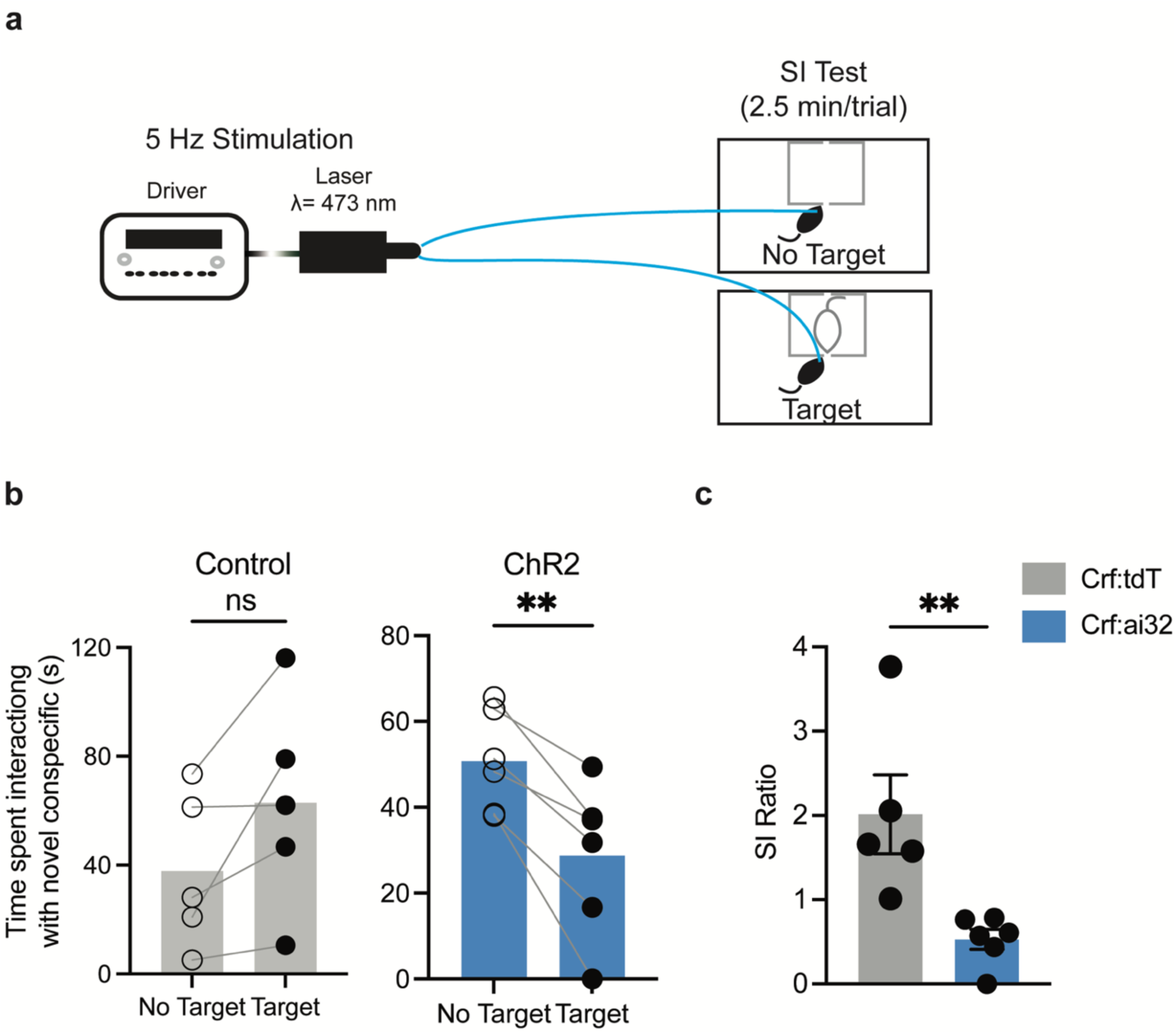
Acute BNSTov^CRF^ stimulation in low- or no-stress conditions induces social avoidance. **a,** 5 Hz optogenetic 473 nm (blue light) photostimulation is induced with 2.5 minutes per trial with no target/target trials in tandem. **b**, Behavior of control mice during social interaction with novel conspecific within two (target vs no target) trials. Paired two-tailed t-test, t=2.282, df=4, P=0.0846, n=5. ChR2 mice receiving photostimulation during social interaction test. Paired two-tailed t-test, t=5.389, df=5, **P=0.0030, n=6. **c**, SI ratio of control and ChR2 mice during the social interaction test. Crf:tdT and crf:ai32, unpaired two-tailed t-test, t=3.352, df=9, **P=0.0085, n=5,6.

Social Interaction (SI) ratio is often used as an index for resilient/susceptible phenotypes based on genetic, electrophysiological, physiological, and behavioral measures.^16–22^ Mice typically prefer interacting with a novel mouse over an empty animal enclosure, as such mice scoring SI ratio>1 are classified as resilient.^16^ Relative to control mice, ChR2 mice showed a significant decrease in SI ratio (Fig 1c). Taken together, this data suggests that in conditions of low stress, BNSTov^CRF^ neurons blunt sociability. Given that in low-stress mice BNSTov^CRF^ neurons promote social avoidance, we then wondered if the stress window between 7 and 10 SDEs is where these neurons acquire the ability to switch from coordinating social avoidance to social approach behavior concomitant with resiliency, in line with previous previous observations.^1^

### Intra-defeat BNSTov^CRF^ neuronal stimulation increases engagement with threatening contexts

Resiliency is associated with the maintenance of spontaneous firing rates in BNSTov^CRF^ neurons between 7 and 10 SDEs. 5 Hz optogenetic stimulation during this stress-sensitive window was sufficient to recapitulate naturally-occurring stress-neuroadaptations that produce resilience. Moreover, cell-type-selective fiber photometry revealed by the 10^th^ social defeat episode (SDE) BNSTov^CRF^ neurons acquire the capacity to respond to social contexts.^1^ However, CRF neural activation does not provide insight into whether social interaction in resilient mice had maintained its expected rewarding properties. In particular, considering the data presented in Fig 1 suggests that BNSTov^CRF^ activation is aversive. Moreover, BNSTov^CRF^ neural activation was shown to promote resiliency, but only when paired with social defeat, suggesting that an essential component of its pro-resilient effect stems from exposure to the stressful context in which maladaptive behaviors could arise. The lines of converging data aforementioned led us to answer several pertinent questions: (1) does stimulation during SDEs 7-10 induce the switch from social avoidance to social approach in resilient mice? (2) how does this intra-defeat stimulation affect social interaction with the familiar CD-1 aggressor present? And (3) does the social behavior toward known aggressors correlate with post-defeat social behavior on the SI test?

During deep brain stimulation (DBS) for treatment-resistant depression, clinicians probe the subject’s emotional state after placing the stimulating electrode. Acute intraoperative stimulation during this placement procedure has been linked to durable DBS antidepressant efficacy.^17^ The solicitation of negative emotions intraoperatively paired with DBS effectively mimics sensitization exposure therapy, where emotional rehabilitation occurs through re-experiencing negative events.^18^ BNSTov^CRF^ stimulation in stress-naïve mice is aversive but produces resiliency in individuals with a history of chronic stress; however, the point at which this transition from aversive to appetitive responses remains elusive. Our previous work revealed BNST stress neuroadaptation occurs between 7 and 10 SDEs is necessary and sufficient for resiliency. Therefore, we hypothesized that 5 Hz photostimulation leverages this window of plasticity to promote resiliency by shifting the valence of BNSTov^CRF^ activation from negative to positive, diminishing the impact of psychosocial stress during repeated social defeat stress (RSDS).

The sensory contact phase is a critical component of RSDS that is necessary to develop susceptibility.^19^ To test this, fiber optic ferrules were implanted over the oval BNST in ai32 transgenic mice crossed with crf-cre lines to allow cre-dependent expression of channelrhodopsin of BNSTov^CRF^ cells. Mice received 5 Hz optogenetic stimulation during the sensory contact phase of the remaining 3 daily SDEs of a 10-day protocol as per previous studies.^1^ We employed behavioral tracking software during optical stimulation to observe the effect of stimulation on social engagement with a familiar CD-1 aggressor (Extended Data Fig 1a,b, Fig 2a,b). Time spent interacting with the CD-1 near the Plexiglas barrier bifurcating the cage were videorecorded, tracked, and scored. CD-1 mice of a larger and more aggressive strain, additionally screened for aggressiveness were used.^16^ The behavioral assay over which experiments took place spanned 8 bins (each bin was 2.5 min). The first bin consisted of non-stimulation-paired social interaction. Bins 2-7 consisted of stimulation, and bin 8 represented a post-stimulation period. On day 1 of stimulation (day 8 of RSDS), there was no significant difference in time spent interacting with the CD-1 aggressor in control mice within, nor across all bins (Extended Data Fig 1c). In contrast, ChR2 mice showed a significant increase in social interaction time across bins compared to control mice (Extended Data Fig 1c,f). On day 2 of stimulation (day 9 of RSDS), there were no significant differences in social investigation between or across bins among control and ChR2 mice (Extended Data Fig 1d,g). On day 3 of stimulation, there was an increase in social interaction of ChR2 relative to control mice when total time is averaged across bins (Extended Data Fig 1e,h). In examining the overall effect of stimulation across all 3 days, it was found that ChR2 mice spent more time on average interacting with CD-1 aggressors at bins 5 and 6 (∼10-12.5 min post-defeat) (Fig 2c). Overall, 5 Hz photostimulation induced significantly higher social interaction times in ChR2 than control mice (Fig 2d). Interestingly, ChR2 mice whose fiber-optic ferrules were implanted off-target were shown to have significantly lower interaction times than both ChR2 and control mice (Fig 2d-f). The stimulation condition in ChR2 mice led to a significant increase in interaction time compared to controls which were not observed in pre- or post-stimulation trials (Fig 2d,e). Among ChR2 mice, interaction time increased during the stimulation phase and persisted post-stimulation (Fig 2e,f). In contrast, no significant differences were observed in the control mice across pre-, stim-and post-stimulation conditions (Fig 2d-f).

**Figure 2.**
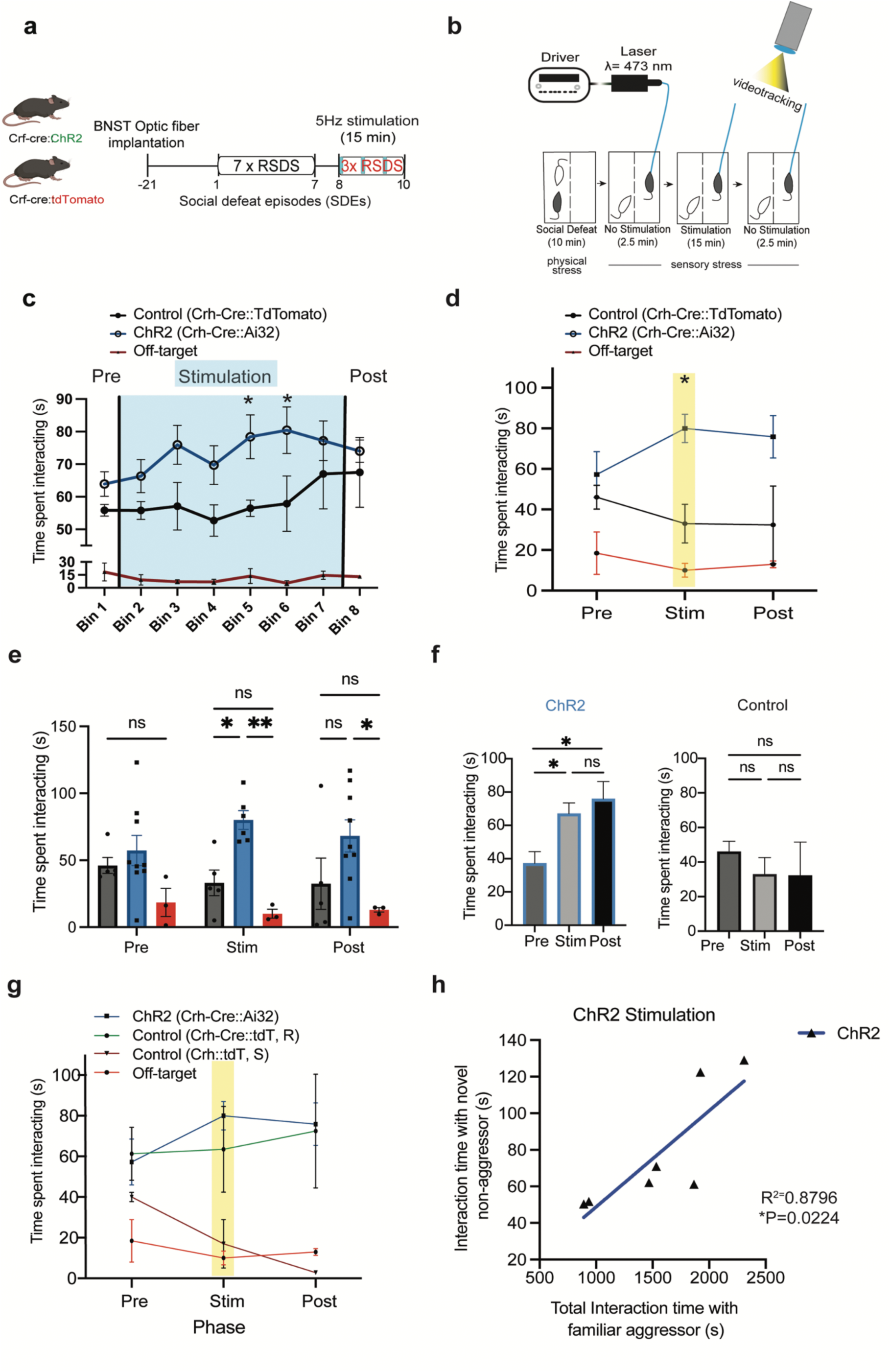
Intra-defeat BNSTov^CRF^ stimulation increases sociability toward familiar CD-1 aggressors and correlates with resiliency development. **a,** Experimental schematic whereby Crf-cre::ChR2 mice are implanted with optical fibers in the BNSTov. **b,** Procedure during SDEs 8 through 10. Mice received optical blue light stimulation for 15 minutes during the sensory stress phase and social interaction is recorded using videotracking software. **c,** Interaction times pre-, intra-, and post-optical stimulation. Control (Crf-cre:tdTomato) n=6; ChR2 (Crf-cre::ChR2) n = 8; Off-target (Crf-cre::ChR2) n = 3. Two-way ANOVA (Bin x stimulation) interaction F(14,47) = 0.6080 P=0.8447, time (Bins) F(7,47) = 0.9344 P=0.4892, stimulation F(2,47) = 208.4 P<0.0001. Tukey’s multiple comparisons post-hoc test, Control vs ChR2 ****P<0.0001, Control vs Off-target ****P<0.0001, ChR2 vs Off-target ****P<0.0001. Bin 5 and 6: Control vs ChR2 *P=0.0438 and *P=0.0369, respectively. Each bin corresponds to 2.5 minutes. **d,** Time spent interacting with all of the bins averaged according to phase. **e,** Time spent interacting across pre-, intra-, and post-stimulation phases. Two-Way ANOVA (Stimulation phase x genotype) F(4,38) = 0.9630, P=0.4389, Stimulation phase F(2,38) = 0.002028 P=0.998, genotype F(2,38) = 16.76, ****P<0.0001. Tukey’s post-hoc test pre-stimulation: Control vs ChR2, Control vs Off-target, ChR2 vs Off-target = P=0.7368, P=0.3446, P=0.0889 respectively. Stimulation: Control vs ChR2, Control vs Off-target, ChR2 vs Off-target = *P=0.017, P=0.475, **P=0.002, respectively. Post-stimulation: Control vs ChR2, Control vs Off-target, ChR2 vs Off-target = *P=0.0192, P=0.5844, **P=0.0037, respectively. **f,** Time spent interacting across SDEs (8^th^-10^th^) according by group. ChR2 (n=8 mice) One-Way ANOVA F(2,44) = 6.118 **P=0.0045 Tukey’s Post-hoc test: Pre vs Stim P=0.0112, Pre vs Post-stim *P=0.0137, Stim vs Post P=0.7521. Control (n=6) One-Way ANOVA Pre vs Stim P=0.7573, Pre vs Post P=0.7362, Stim vs Post P=0.9993. **g,** Retrospective analysis of stimulation during SDEs 8-10 effect on susceptible/resilient mice. ChR2 (n=8 mice), Control (R) (n= 2 mice), Control (S) (n = 4 mice), Off-target implanted ChR2 (n = 3 mice). Two-Way ANOVA Interaction (Stimulation phase x Condition) F(6,38) = 0.6668 P=0.6768, stimulation phase F(2,38) = 0.03819 P=0.9626, condition F(3,38) = 10.86 ****P<0.0001. Tukey’s post-hoc test: stimulation ChR2 vs Off-target *P=0.0114. Post-stimulation: Control (R) vs Control (S) = P=0.0521, ChR2 vs Off-Target *P=0.019, ChR2 vs Control (S) *P=0.02. **h,** Correlation of intra-defeat stimulation (familiar aggressor) by social interaction time on the SI test (novel non-aggressor conspecific). ChR2 (n=9), linear regression goodness of fit test, ChR2 R^2^=0.4979, F(1,7) = 6.941, *P=0.0337.

Per established protocols, c57bl/6j mice undergoing 10 days of RSDS will phenotypically display susceptibility or resiliency; therefore, the control group’s mean masks the effects of these subgroups. Interestingly, when the control group is parsed into their respective phenotypes (based on SI ratio^16, 20^, *see methods*), susceptible mice were observed to have similar interaction times as the off-target ChR2 group (Fig 2g). Conversely, resilient mice displayed similar social interaction times as the ChR2 group (Fig 2g). The regression of intra-stimulation interaction time with CD-1 aggressor and time spent with a novel conspecific at a later social interaction test revealed a positive and significant correlation in ChR2 but not control mice (Fig 2h, Extended Fig 1i). Optogenetic stimulation appeared to prevent the eventual decline in of social interaction time with familiar aggressors in ChR2 that occurred in control mice amidst repeated attacks, thought this effect was not statistically significant (P=0.0524, Extended Data Fig 1j). In summary, these results reveal that neuromodulation during this stress window is crucial to the development of resiliency because it provokes desensitization exposure, which may neutralize the emotional valence of psychosocial aggression.

### Optical real-time place preference reveals that BNSTov^CRF^ neuronal modulation biases social preference

While BNSTov^CRF^ neurons robustly produce resilient phenotypes, it is unclear what motivates the observed increase in social interaction. On the one hand, increased pro-social behavior could be due to stress-related hypervigilance governed by a negative internal state because of cumulative stress exposure. Conversely, increased sociability could result from an enhancement in the rewarding properties of social contact. The distinction has clinical implications as many stress-related psychopathological states manifest as “functional” behaviors, such as feigned sociality, despite distressing internal states generated by such behaviors.^10, 21, 22^ Whereas, clinically, resilience represents an attenuation of negative affect, giving rise to the internal experience of rewarding social interactions. Therefore, we tested whether a positively motivated state drives the social behavior that characterizes resiliency. To answer this question, we used a modified social optical-real time place preference assay, or social o-RTPP, whereby a mouse was permitted to navigate between two compartments containing novel non-aggressive male CD-1s, with one paired with optical stimulation. Time spent in each chamber was used to determine the effect of stimulation on preference. Control mice expressed no preference, as the percentage of time spent was similar across both chambers (Fig 3a). In contrast, ChR2 mice demonstrated a strong preference for the photostimulation paired chamber (Fig 3b), as evidenced by the increased percentage of time spent in the chamber, specifically when co-paired with stimulation (Fig 3c). Spending more time in a compartment demonstrates side preference but may not represent social preference, as spatial tracking does not consider specific actions undertaken in the chambers that may confound results such as distinguishing freezing versus interactions with the CD-1. Therefore, we analyzed, posthoc, the percentage of time ChR2 mice spent explicitly interacting with a novel conspecific both in the presence and absence of photostimulation. We uncovered that mice spent three-fold more time engaged in social interaction with the novel CD-1 while in the preferred chamber compared to the unpaired trial (19.80+/-9.096 vs. 59.40+/-4.167, no light vs. light, respectively, Fig 3d). These results suggest BNSTov^CRF^ stress neuroadaptation coordinates resilient behavioral responses through generating appetitive, pro-affiliative social preference.

**Figure 3.**
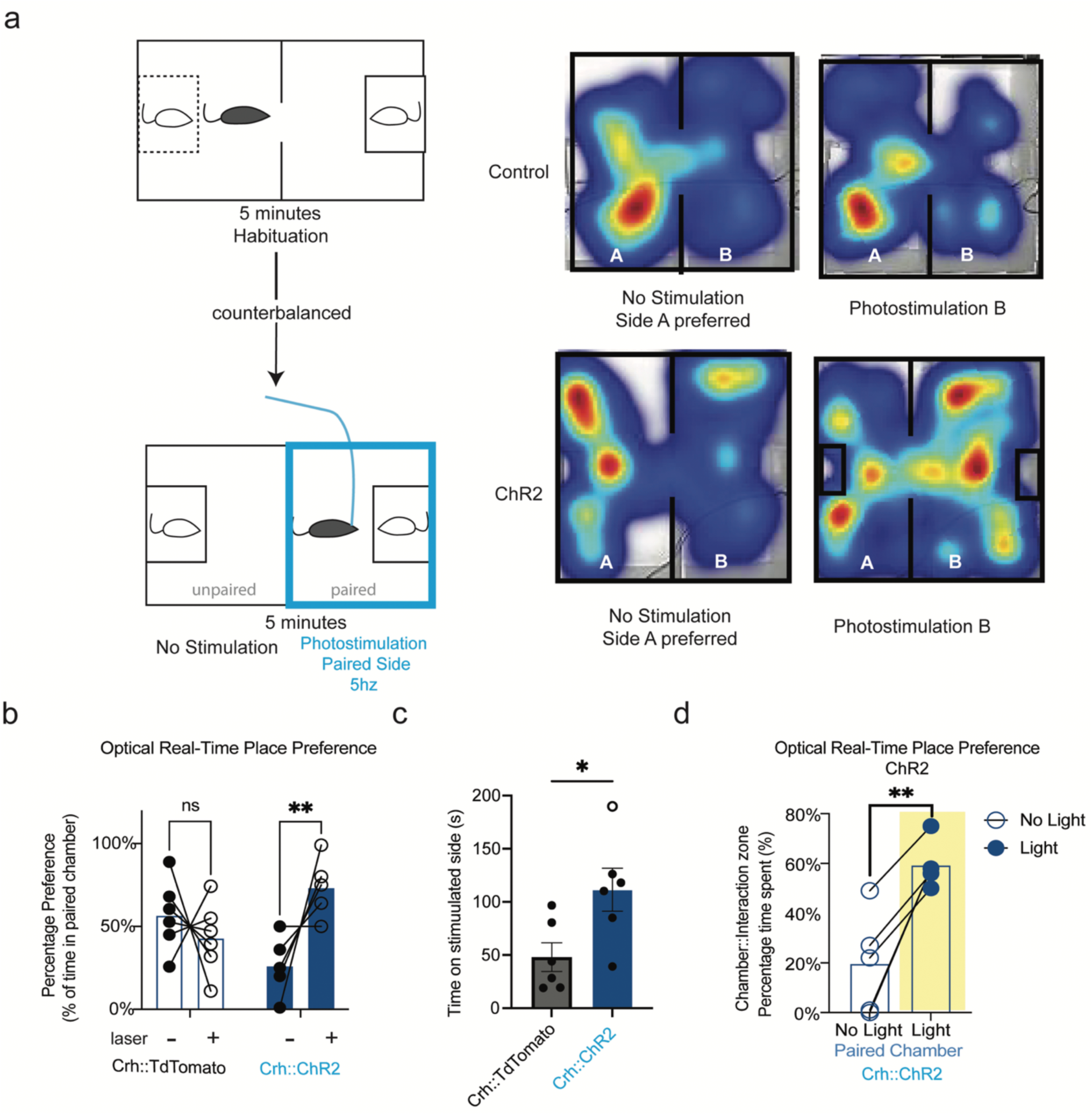
BNSTov^CRF^ neurons shift socio-affective bias on social optical real-time place preference assay. **a,** Schematic of social optical real-time place preference (o-RTPP), where mice have a 5 minute habituation followed by photostimulation paired to the side mice where spent the least amount of time during habituation. This is reflected in the heatmap of control and ChR2 mice. **b,** Social o-RTPP, where control crf:tdTomato and Crf::ChR2 mice spend more time on the laser paired side only in the ChR2 mice. Control = 5 mice, ChR2 = 6 mice. Two-way ANOVA Interaction (laser_on/off_ x genotype_control/ChR2_) F(1,18) = 0.0023 **P=0.0023. Genotype F(1,18) = 0.0000 P>0.9999. Laser F(1,18) = 3.792 P=0.0673. Sidak’s post-hoc test Control P=0.4418, ChR2 **P=0.0032. **c,** Time spent on stimulated side between control and ChR2 mice. Unpaired t-test, two-wailed t=2.603, df=10, P=0.0263. **d,** Social o-RTPP of ChR2 mice as a percentage of time spent interacting when in stimulation paired chamber. Stimulation specially induces social interactivity when mice prefer a side (n=5 mice). Paired t-test, two-tiled, d=5.406, df=4, **P=0.0057.

### BNSTov^CRF^ stress neuromodulation enhances positive valence to appetitive and aversive socially-salient stimuli

Resiliency involves gathering information about environments containing a variety of emotionally-valenced stimuli ranging from reward-to threat-related cues for survival.^10, 23, 24^ Rodents navigate using olfactory cues, and the BNST receives direct input from the vomeronasal accessory olfactory system.^25, 26^ Stress modifies BNSTov^CRF^ neurons in resilient mice to maintain adaptive social behaviors in social defeat stress (Extended data Fig1c-g), but we wondered whether this could generalize across contexts of varying valence. To test this, mice were placed in a three-chamber arena over several trials that contained distinct odorants of varying valence: water, male urine, female urine, and predator (Red fox) urine (Fig 4a, Extended Data Fig 2a) and measured investigation times in trials lasting two-minute trials of 5 Hz stimulation (trial 1: “light off”, trial 2: “light on”). There was no significant difference in time investigating water between ChR2 and control mice (Fig 4b). Similarly, ChR2 and control mice expressed no difference in the investigation of male urine (Fig 4c). ChR2 compared to control mice spent significantly more time investigating female urine, suggesting that BNSTov^CRF^ enhances the valence of rewarding stimuli (Fig 4d). Indeed, resilient mice spent more time investigating female urine during the no stimulation trial (Extended data Fig 2b). To examine BNSTov^CRF^ neurons’ role in socially salient aversive stimuli, mice were subjected to predator urine. Predator urine odorant has been described as an innate fear stimulant,^27, 28^ and lesions of the dorsolateral BNST (a region that includes the oval nucleus) inhibit defensive/freezing responses in its presence. Interestingly, we observed that BNSTov^CRF^ activation promoted increased investigation and sniffing of predator urine compared to control mice (Fig 4e). This finding suggests that BNSTov^CRF^ neurons promote resiliency, in part, by shifting the valence of aversive contexts in response to stress. Notably, in the absence of optical stimulation (“no light” trials), there was no significant difference in sniffing time among all odorants between ChR2 and control mice (Fig 4f). Moreover, there was no significant difference between susceptible/resilient mice in investigation male urine during the no-light trials (Extended data Fig 2b). Predator urine solicited no significant differences between susceptible/resilient mice reflecting the innate fear potency of the odorant, as both groups investigated the stimulus for a very short period (Extended data Fig 2b). In summary, these findings suggest that resiliency induced by photostimulation between the 7th and 10^th^ SDE promotes the investigation of a range of socially derived stimuli essential to survival.

**Figure 4.**
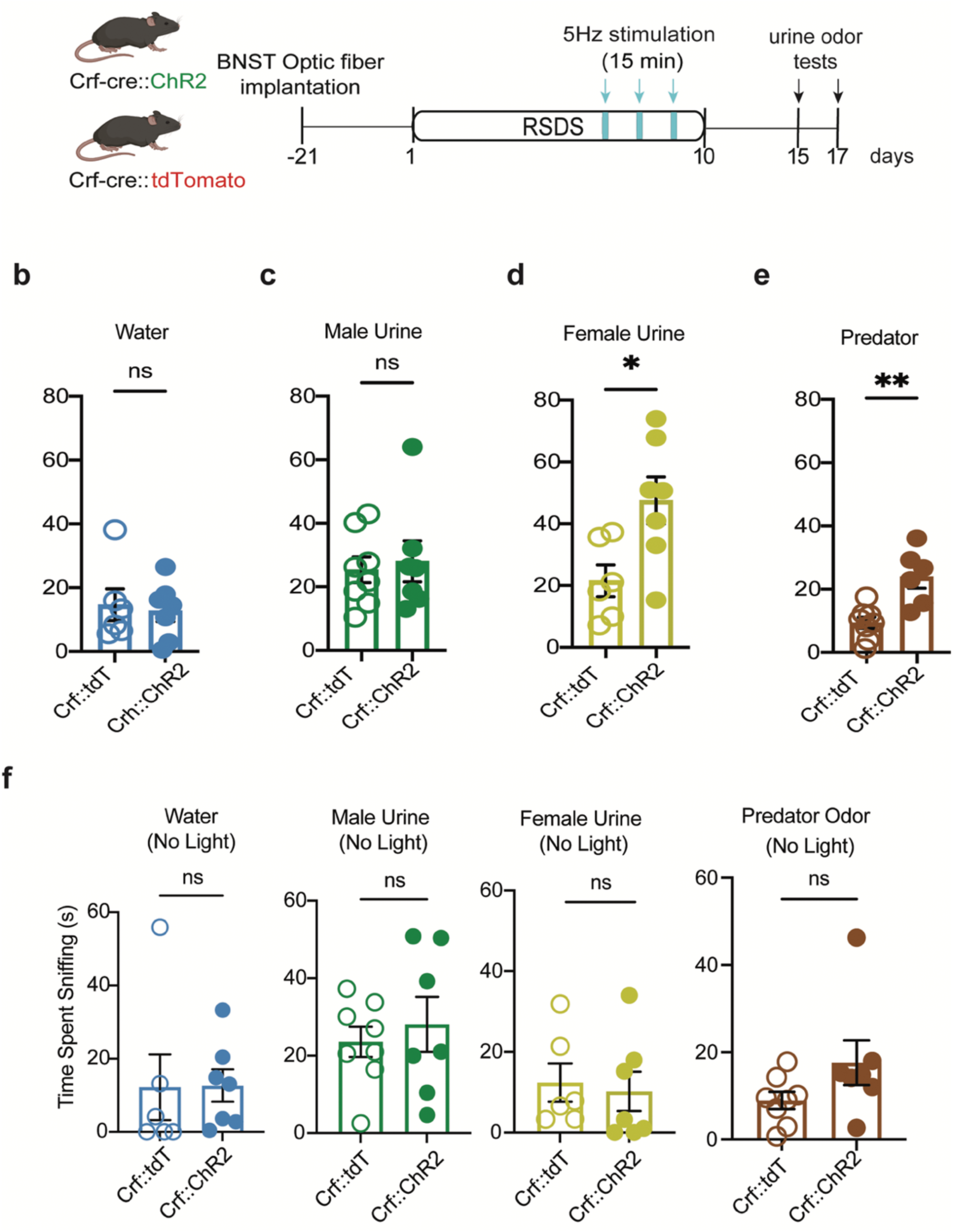
BNSTov^CRF^ stress modulation enhances positive valence to appetitive and aversive socially-salient stimuli. **a,** Experimental timeline, pending fiber optic implant mice are subjected to 10 SDEs with concomitant optogentic stimulation the final three episodes of RSDS and exposed to water, male-, female-, and predator urine over two days. **b,** The time spent sniffing water was recorded on behavioral recording software and quantified under direct effect of optical stimulation. Crf::tdt = 6 mice, Crf::ChR2 = 7 mice, unpaired two-tailed t-test, t=1.561, df=11 P=0.1469. **c,** The time spent sniffing male urine was recorded on behavioral recording software and quantified under direct effect of optical stimulation. Crf::tdt = 8 mice, Crf::ChR2 = 7 mice, unpaired two-tailed t-test, t=0.3592, df=13 P=0.7252. **d,** The time spent sniffing female urine was recorded on behavioral recording software and quantified under direct effect of optical stimulation. Crf::tdt = 6 mice, Crf::ChR2 = 7 mice, unpaired two-tailed t-test, t=2.726, df=11 *P=0.0197. **e,** The time spent sniffing predator urine was recorded on behavioral recording software and quantified. Crf::tdt = 8 mice, Crf::ChR2 = 6 mice, unpaired two-tailed t-test, t=3.943, df=12 **P=0.002. **f,** The time spent sniffing male urine was recorded on behavioral recording software and quantified in the absence of optical stimulation. Unpaired two-tailed t-test, water: t=1.561, df=11, P=0.1469; male urine: t=0.3592, df=13 P=0.7252; female urine: t=1.561, df=11 P=0.1469; predator urine: t=1.661, df=13 P=0.1207.

### BNSTov^CRF^ neuronal activation suppresses despair-like behavior on the tail suspension test

BNSTov^CRF^ neuronal activation promoted increased investigation of a familiar aggressor CD-1 (Fig 2d) and predator odorant (Fig 4e), suggesting that this increased activation may promote an increased tolerance for stressors. However, in both, mice had control over their proximity to the potentially threatening stimuli. Therefore, it challenges interpretations of whether activation of BNSTov^CRF^ neurons in an inescapable non-social physical stressor would promote resiliency. To test this, we utilized the tail-suspension test (TST) with optogenetic activation following RSDS and photostimulation during 8-10 SDEs (Fig 5a). Mice were initially suspended without stimulation for the first three minutes, and no significant differences between control and ChR2 mice were observed (Fig 5b). During the three-minute period of stimulation, ChR2 mice spent less time immobile than control mice (Fig 5b). Intriguingly, in the final three minutes following stimulation, ChR2 spent less time immobile, suggesting the internal state induced by stimulation persisted (Fig 5c). In summary, we show that BNSTov^CRF^ activation in a variety of stress-related contexts causes a positive shift in affective state, thereby motivating behaviors consistent with resiliency.

**Figure 5.**
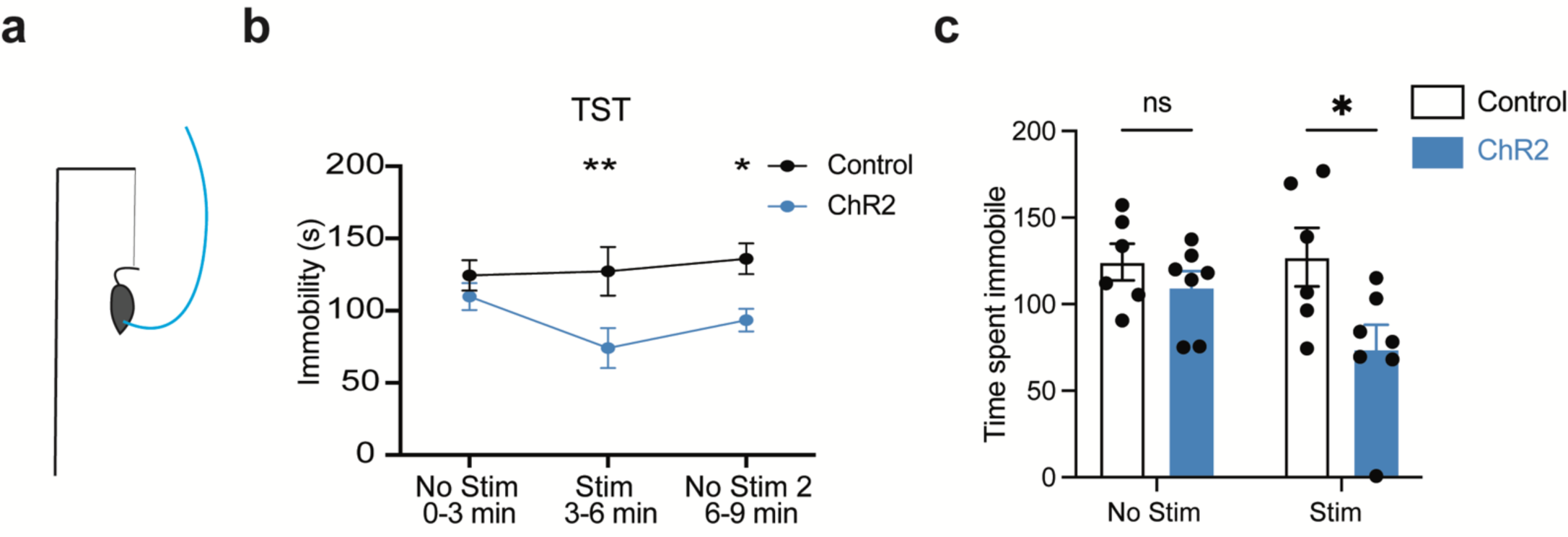
Optogenetic activation of BNSTov^CRF^ neurons suppresses despair-like behavior on TST that outlasts acute stimulation. **a,** Tail suspension test scheme, whereby mice were suspended by tail and were tethered to an optogenetic fiber optic patch cord. **b,** Time spent immobile on the TST during no stimulation, stimulation, and post-stimulation phases, lasting 3 minutes each. Control = 6 mice, ChR2 = 7 mice. Two-Way ANOVA Interaction (Stimulation phase x Condition(genotype)) F(2,33) = 1.410 P=0.2584, Stimulation phase F(2,33) = 1.128 P=0.3357, Condition F(1,33) = 14.53 ***P=0.0006. Sidak’s Post-hoc test: Control – ChR2 (no stim): P=0.768, (stim) **P=0.0096, (no stim 2) *P=0.0467. **c,** Time spent immobile on TST as a feature of control vs ChR2 conditions. Two-Way ANOVA Interaction (Stimulation phase x Condition(genotype)) F(1,22) = 2.209 P=0.1514, Stimulation phase F(1,22) = 1.616 P=0.2169, Condition F(1,22) = 6.881 *P=0.0155. Sidak’s Post-hoc test: No Stim – Stim (control): P=0.6752, (ChR2) *P=0.0163 Extended Data Fig 1: Intra-defeat stimulation increases interaction time with familiar CD-1 aggressors in ChR2 not control mice.

## Discussion

By capturing direct social behavioral effects of 5 Hz optical stimulation, we observed increased sociability toward a familiar aggressor, an experience crucial to establishing resiliency. Photostimulation of stress-naïve and singly defeated mice induced social avoidance, reflecting the baseline role BNSTov^CRF^ neurons play in social contexts. Using social o-RTPP, we observed the effect of BNSTov^CRF^ activation on shifting socio-affective bias. This activation also modulated salience of odor-based, socially-valenced stimuli, as optogenetic stimulation-induced increased investigation of female urine and surprisingly predator urine, an innate stressor. Lastly, TST was used to assess whether the resiliency-inducing stimulation protocol had influenced adaptive responding to inescapable, non-social stressors. We uncovered that photoactivation induced a decrease in immobility that persisted after stimulation. The experiments together provide important insight into how BNSTov^CRF^ neurons induce resilience by maintaining a positive internal state resistant to the insults of ongoing stress.

In this report, we observe that stress modulation of the BNSTov produces resilience by a positive shift in affective state. Unmitigated stress exposure has a deleterious effect on emotional state producing despair-like behavior and learned helplessness.^29–31^ We predicted that CRF activation would promote the internal states that produce despair-like behavior. Surprisingly, in a similar manner to our previous work on resiliency, we find that post-stress modifications to BNSTov^CRF^ neurons favor a positive internal state in the face of ongoing social stress. We observed that intra-defeat optogenetic stimulation during 8-10 SDEs, promoted more social interaction with a familiar aggressor. We show a significant positive correlation between interaction time with the aggressor and sociability to novel non-aggressive mice following 10 defeat episodes.

The sensory contact phase of RSDS is critical to the phenotypic display of resilience/susceptibility, as elimination of this component of RSDS failed to produce susceptible mice.^19, 32^ Therefore, the 15 minutes of sensory stress on 8^th^-10^th^ SDEs may represent a critical time during which emotional learning occurs. The development of resiliency may be shaped by emotional learning that occurs in the context of experiencing cumulative stress insults.^9, 33^ Indeed, mice implanted with fiber-optic ferules off-target failed to become resilient, suggesting that BNSTov^CRF^ engagement is protective during RSDS and the absence of which leads to susceptibility. Similarly, control mice (effectively serving as a sham group) did not display resiliency, further illustrating the importance of BNSTov^CRF^ activation for emotional learning. Male mice are innately territorial and engage in agonistic interactions with other mice, even in laboratory settings.^34 35^ Given the natural tendency for mice to defend their territory, repeated attacks from a larger mouse of a more aggressive strain would constitute a form of social conditioning.^36^ During each SDE, mice acquire more information about their subordinate status where they assume behavioral changes, such as defensive posturing, consistent with susceptibility.^36^ This form of emotional learning is referred to as conditioned defeat.^36–39^ The BNST has been shown to be necessary for conditioned defeat status.^27^ Therefore, BNSTov^CRF^ stimulation during the period between 8 and 10 SDEs likely impaired the internal aversiveness of being defeated, impacting the way defeated mice would encode the experience. The stimulation-induced experience may dampen social defeat’s aversiveness enough that conditioned defeat does not occur. In this study, optogenetic stimulation increased the time spent investigating CD-1, likely indicating the optical stimulation changed the subjective experience of stress, as male mice would typically avoid such a conspecific. Another plausible explanation is that stimulation increased vigilance, as BNST activation has been associated with increased arousal and anticipatory anxiety.^40, 41^ However, the effect of stress on motivated social behavior could also be due to a blunting of circuits coordinated through the BNST in susceptible mice.

A possible explanation for why intra-defeat sociability predicts post-defeat resiliency is that the period comprising 8-10 SDEs represents a unique window by which mice assess their controllability, in order to maximize behavioral adaptation to this setting. Photostimulation may catalyze the continual promotion of behavioral flexibility and active coping by shifting internal states.^11, 42, 43^ Stimulation induces plasticity that allows incorporating a more optimistic context for a stressful environment. Cognitive tasks using instrumental conditioning, for instance, would be needed to assess this adequately. In fact, leveraging plasticity for therapeutic efficacy is shown in recent reports to play a role in the success of deep brain stimulation (DBS) for treatment-resistant depression. A recent report showed that individualized tractography-guided implantation and intraoperative stimulation predicted early antidepressant effects ^17^. Previous work manipulated BNSTov^CRF^ neurons in the social defeat context, yet the behavioral effect was monitored in a separate novel context, only allowing inference of the direct stimulation on social behavior. Using social o-RTPP, we assessed the effects of neuronal stimulation on bivalently coordinating the internal emotional state. We observed that resiliency could effectuate both a state and trait, as 89% of the ChR2 mice were resilient and showed a preference for the side and social target in which they were stimulated, suggesting a state-dependent enhancement. In the context of socially-salient stimuli, such as female odor exposure, resilient mice displayed increased time investigating compared to susceptible mice, which was further enhanced during the “stimulation on” trials. This finding underlies how resilient mice may come to display higher social interaction times and hedonic-like consumption following BNSTov^CRF^ stimulation because it induces a positive internal state that enhances experiences that are generally positive. Notably, this effect is not simply relegated to all stimuli, as water or male urine did not solicit increased time sniffing during photostimulation. However, we cannot discount the modulatory role BNSTov^CRF^ neurons play in producing susceptibility. Indeed, in mice, we see the neuronal adaptation in both electrophysiological measurements and in calcium dynamics that support a state of decreased activation, suggesting that resiliency is the maintenance of activity in the face of threatening stimuli. This adaptation explains why photoactivation increased time spent sniffing predator urine, an innate stressor. Freezing to predator urine is thought to be BNST dependent, as lesions reduce defensive responses by way of the vomeronasal system. Interestingly, BNSTov^CRF^ activation would be predicted to increase freezing by synergizing accessory olfactory-BNST circuits^44–46^ instead, we observed the opposite. This interpretation is reinforced by the effects of BNSTov^CRF^ stimulation during TST in decreased immobility, suggesting that resiliency maintains positive internal states in the face of threats by neuronal activation.

A question that emerges from the data is, whether it is advantageous for an organism to frame aversive experiences more positively? After all, decreased exposure to known threats such as a familiar aggressor or predator urine has survival implications.^44, 47^ On the other hand, the role of BNSTov^CRF^ in valence switching promotes cognitive flexibility, decision-making, and social coordination, which can assist in foraging limited resources.^46, 48–53^ The BNST is important for integrating homeostatic needs with external input.^42^ Mice in the study were sated and had *ad libitum* access to water; it would be interesting to see how behavioral responses altered to meet homeostatic demands.

An important caveat to this study was that mice were subjected to intra-defeat optical stimulation, which produced resiliency, and later placed in a variety of behavioral tasks for which acute optogenetic activation occurred. A strength of this study design is that it enables the assessment of resiliency as a trait vs. state. Traits represent a stable and enduring pattern of behavior, whereas a state is a temporary mode of being relegated to acute circumstances. Our previous work focuses on resiliency as a trait, as mice displayed hedonic-like and pro-social behavior lasting up to 6 weeks^1^. In this report, we uncover that acute optogenetic activation of (optically-induced) resilient mice enhances motivated exploratory behavior as a state. State-dependent resiliency is evident in the urine test, where no-light trials yielded no difference between control and ChR2 mice. In contrast, photostimulated ChR2 mice promoted increased female urine sniffing and social interaction with a novel conspecific, both of which are already elevated in resilient mice in the absence of manipulation.

Additionally, inhibitory optogenetics could explore the necessity and sufficiency of BNSTov^CRF^ neurons to state-dependent behaviors. Future studies could parse the contribution of state vs. trait by optogenetically activating during mice during behavioral tasks only after 10 SDEs. As neuromodulation becomes a more favored approach to treating mood disorders, an appreciation for stress-dependent changes in internal state may favor stimulation protocols (for DBS or transcranial magnetic stimulation) that may work in one context and not another. Thus, pioneering closed-loop or neurofeedback components into the stimulating device cannot be untethered from the patient’s emotional reports, as they may serve as proxies for neurophysiological adaptations pertinent to antidepressant efficacy.

## Methods

### Mice

The study used wild-type, Crf-ires-Cre (Jackson labs: 011087), ai14 (Cre-responsive tdTomato reporter mouse; Jackson labs: 007915), ai32 (Cre-responsive channelrhodopsin-2/fused with eYFP; Jackson labs: 012569) mice on C57BL/6J background that were bred at Icahn School of Medicine at Mount Sinai and used were between 6-7 weeks at the start of experimental manipulations. Upon receipt from Jackson Laboratories, mice were acclimated to the housing facility for 1-2 weeks prior to the start of experiments. All mice were group-housed and maintained on a 12/12 light-dark cycle with ad libitum access to food and water. Following social defeat stress, mice were singly housed and maintained on a 12/12 light-dark cycle with ad libitum access to food and water. All experiments were approved by the Institutional Animal Care and Use Committee and comply with institutional guidelines for the Animal Care and Use Committee set forth by Icahn School of Medicine at Mount Sinai.

### Repeated social defeat stress paradigm

The repeated social defeat stress paradigm was performed according to published protocols^16, 20, 54–58^. Briefly, CD1 aggressors were singly housed in cages (26.7 cm width x 48.3 cm depth x 15.2 cm height; Allentown Inc) at least 24 hours before the start of the experiment on one side with a clear perforated Plexiglas divider. Social stress consists of physical and sensory stress components. During the period which marks the physical stress, C57BL/6J mice are placed on the ipsilateral side of the cage as the CD1 aggressor for 10 minutes. Following this, the intruder mouse is placed in the contralateral side of the perforated Plexiglas divider for the remainder of the 24-hour period, marking the sensory-stress period. Every 24 hours, the intruder mouse is paired with a new aggressor for 10 episodes. Control mice were housed two mice per cage divided by a perforated Plexiglas divider and rotated and handled daily like the socially defeated mice.

### Social interaction test

Social interaction testing was performed as described^16, 20, 23, 58–62^. Briefly, a novel conspecific of CD1 strain is placed in an interaction zone of a standard open-field arena, and the time the intruder spends in the interaction zone is measured. The mice spend a total of 5 minutes in the open arena (2.5 min with and without the novel CD1). Ethovision (Noldus Information Technology) video-tracking software is used to track interaction time. All social interaction testing takes place 24 hours after the last defeat. Social interaction (SI) is measured by time spent in the interaction zone during first (CD1 absent) over second (CD1 present) trials. Mice are categorized according to SI ratios; an SI ratio ≧1 defines resilient, whereas an SI ratio <1 is susceptible as described previously^9, 20, 63, 64^.

### Sucrose preference test

For sucrose preference testing, a solution of 1% sucrose or diluent alone (drinking water) is filled in 50 ml tubes with ball-pointed sipper nozzles (Ancare). Animals are acclimatized to two-bottle choice conditions prior to testing. The bottles are weighed, and positions interchanged daily. Sucrose preference is calculated as a percentage [100 x volume of sucrose consumed (in bottle A)/total volume consumed (bottles A and B)] and averaged over 2 days of testing.

### Stereotaxic virus and optic fiber implantation

Under ketamine (80 mg/kg)/xylazine (10 mg/kg) anesthesia, mice were placed in a stereotaxic frame (Kopf Instruments) and the BNSTov was targeted (coordinates: anterior/posterior +0.20, media/lateral: +/-2.15, dorsal/ventral, –4.0 mm; 15-degree angle). Hair was shaved around the crown of the head, alcohol, and betadine were applied to the scalp. Ophthalmic ointment was applied to the eyes to prevent dryness, a midline incision was made down the scalp, and a craniotomy was made using a dental drill. A 10 ul Nanofil Hamilton syringe (WPI, Sarasota, Fl) with a 34-gauge beveled metal needle was used to infuse 0.5 ul virus at a rate of 85 nl/minute. Following infusion, the needle was kept at the injection site for 10 minutes and slowly withdrawn. For the optogenetic experiments, chronically implantable optic fibers constructed with 1.25 mm diameter, 200 um core, 0.39 numeral aperture (NA) were used (RWD). For the fiber photometry experiments, chronically implantable optic fibers constructed with 1.25 mm diameter, 400 um core 0.48 numerical aperture (NA) optic fiber and unilaterally implanted into the BNSTov (coordinates: anterior/posterior +0.20, media/lateral: +/-2.15, dorsal/ventral, –3.9 mm; 15° angle) (thor labs). Fiber optical ferrules were cemented to the skull using dental acrylic (Parkell C&B Metabond). All optical stimulation and fiber photometric recording experiments were conducted a minimum of 3-4 weeks post-implantation.

### Optogenetic manipulation of BNSTov^CRF^ neurons

Optical fibers were implanted on Ai32 (transgenic mouse line expressing light-gated cation channel channelrhodopsin-2 (ChR2). Optical fiber Optogenetic stimulation was conducted via the use of a diode-pumped solid-state (DPSS) 473-nm blue laser (Crystal Laser, BCL-473-050-M), using a patch cord with an FC/PC adaptor (Doric Lenses, MFP_200/240/900-0.22_4m_FC-MF2.5). A functional generator (Agilent Technologies; 33220A) was used to generate a 5 Hz frequency, pulse width of 10 ms for 15 minutes^19, 32^, and power density between 7-9 mW mm^-2.^ Experimenters were blinded to the stimulation group.

### Social Optical Real-Time Place Preference

In this test of sociability, mice were placed in a 3-chamber rectangular apparatus (61 cm x 40.5 cm x 23.5 cm) with clear acrylic walls and a white matte flooring. Initially, mice are habituated by being placed inside the center chamber for 5 minutes after being tethered to fiber optic cables. The second phase of 5 minutes of the experiment, occurs with the placement with the placement of 2 CD-1 male mice that were placed in target enclosures with grid-iron opening on one side to enable social interaction. The CD-1 mice were used only if they consistently failed screening criteria for being use in social defeat stress, and in a separate social interaction test mice display social interest and zero antagonistic behaviors. During the 5 minutes of phase 2 mice are tracked using Ethovision 10 (Noldus) and separate time spent in chamber side and interaction time were collected. Data from phase 2 were assessed “on-line” directly afterward and a side preference was identified. All mice showed a preference for one CD-1 compared to the other. On the third phase of the experiment (5 minutes), 5 Hz stimulation was assigned to the chamber opposite of that which the mouse preferred in phase 2. During the stimulation session, optical stimulate was delivered whenever the mouse enters into the stimulation chamber and was stopped once the mouse left.

### Salient Urine Odorant Exposure

Odorants were placed in a 3-chamber rectangular apparatus (61 cm x 40.5 cm x 23.5 cm) and water, female, male, or fox urine was placed on 1 cm x 1 cm pieces of cotton (Nestlet). and contained in 2 cm diameter miniature petri dishes. Mouse urine and water volumbers were 500 µl, fox urine was 5 µl. Mice were tested over 3 trials over 3 days, each trial consisted of 2 minutes. Trial 1 was water vs male urine, trial 2 was male urine vs female urine, trial 3 was male urine vs predator urine and time spent interacting between the odorants were tracked with ethovision (Nodus). Odorants were counter-balanced on sides across mice and conditions (control vs ChR2 mice). Sniffing was described as nose to nestlet.

### Immunohistochemistry

Mice were quickly anesthetized with urethane and perfused with cold 1x-PBS (OmniPure, Bio-Rad) followed by cold 4% PFA (Fisher Hamilton Scientific). Brains were placed in PFA at 4°C for at least 24 hours, followed by being replaced with a 15% sucrose/PBS solution at 4°C for 24 hours and brain sinking. The next day, brains were placed in a 30% sucrose/PBS solution at 4°C for 24 hours. Once the brains sunk, it was washed in 1xPBS three times and mounted on a freezing microtome for sectioning at 35 micrometers.

Sections were washed three times for 10 minutes in 1x PBS, then blocked in 5% bovine serum albumin (Sigma-Aldrich) in 1x PBS with 0.3% Triton-X (Sigma-Aldrich) for 1 hour. Tissues were incubated with primary antibodies for rabbit anti-mCherry ((1:1000), Invitrogen), rabbit anti-c-Fos antibody (1:250, Milipore C-10, sc-271243) and goat anti-GFP (1:500, abcams) overnight at 4°C. Sections were washed three times with 1xPBS and blocked with 0.3% Triton-X for one hour. Tissue was blocked with secondary antibodies goat anti-rabbit/568 (1:1000, Abcams) and donkey anti-goat Alexa-488 (1:500, Abcams) at room temperature for 1 hour. Sections were washed three times with 1x PBS for 10 minutes and mounted on slides with ProLong Gold antifade reagent with DAPI (invitrogen, P36931). Z-stacked images were acquired with a Zeiss LSM780 multi photon confocal system and z-stacks were collated using ImageJ (Fiji). Cell counters were obtained using the Cell Counter feature on ImageJ.

### Statistical analysis

Animals were randomly assigned to control and experimental groups and all experimenters were blinded. Mice were excluded if viral infection was off-target. No data was excluded for other reasons. Student’s two-tailed t-tests were used for comparisons of two experimental groups. For parametric data sets, comparisons among three or more groups were performed using one or two-way ANOVA tests followed by Tukey’s or Bonferroni posthoc tests. For all tests, p<0.05 was determined to be significant. Statistical analyses were performed using Graph Pad Prism 9 .3.1 software (La Jolla, CA, USA). For data not normally distributed, non-parametric analyses were performed.

## Acknowledgements

We would like to thank S.M. for generous sharing for input on the manuscript. We would like to thank S.R. and L.L. for lending viral constructs for behavioral validation. We would like to thank the animal care staff at ISMMS for the animal care staff and technical support. This research was funded by the National Institutes of Health grants R01MH072908 to DGR and LJY, R01MH1206387 to MHH and SJR, R21MH112081 to MHH and SJR, F31MH114624 to SEH, and P51 OD011132 to YNPRC, by the National Key R&D Program of 1034 China No. 2021ZD0202900 to MHH. Figure 1. Acute BNSTov^CRF^ stimulation in low- or no-stress conditions induces social avoidance.

---

**Extended Data Fig 1:**
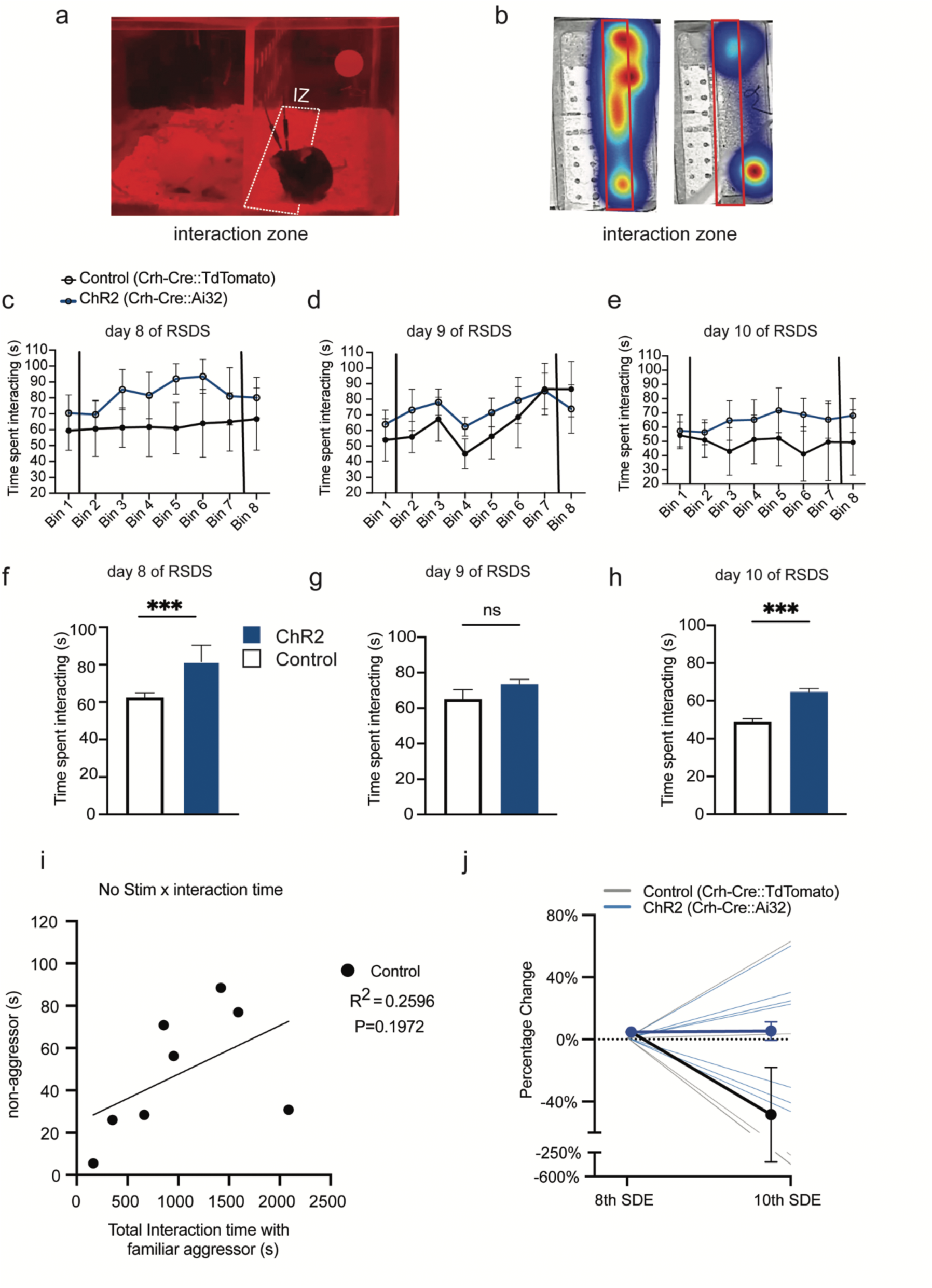
Intra-defeat stimulation increases interaction time with familiar CD-1 aggressors in ChR2 not control mice. **a,** Day 8 of RSDS interaction times, vertical bar separate pre-, intra-, and post-optical stimulation phases, respectively. Control (Crf-cre:tdTomato) n=6; ChR2 (Crf-cre::ChR2) n = 8; Two-way ANOVA (Bin x stimulation) interaction F(8,105) = 0.2587 P=0.9775, time (Bins) F(8,105) = 0.7179 P=0.6753, stimulation F(1,105) = 6.175 *P<0.0145. Each bin corresponds to 2.5 minutes. **b,** Day 9 of RSDS interaction times, vertical bar separate pre-, intra-, and post-optical stimulation phases, respectively. Control (Crf-cre:tdTomato) n=6; ChR2 (Crf-cre::ChR2) n = 8; Two-way ANOVA (Bin x stimulation) interaction F(7,103) = 0.3130 P=0.9467, time (Bins) F(7,103) = 1.356 P=0.2318, stimulation F(7,103) = 1.662 P=0.2003. Each bin corresponds to 2.5 minutes. **c,** Day 10 of RSDS interaction times, vertical bar separate pre-, intra-, and post-optical stimulation phases, respectively. Control (Crf-cre:tdTomato) n=6; ChR2 (Crf-cre::ChR2) n = 8; Two-way ANOVA (Bin x stimulation) interaction F(7,99) = 0.1448 P=0.9943, time (Bins) F(7,99) = 0.6926 P=0.9995, stimulation F(1,99) = 4.228 P=0.0424. Each bin corresponds to 2.5 minutes. **d**, Interaction time with familiar aggressor during 15 minute stimulation averaged across bins on day 8 of RSDS. Two-tailed Mann-Whitney T-test **P=0.0040, (66.28 +/-3.918, control) vs (83.06 +/-3.068 ChR2), n=9,9. **e,** Interaction time with familiar aggressor during 15 minute stimulation averaged across bins on day 9 of RSDS. Two-tailed Mann-Whitney T-test P=0.2345, (65.00 +/- 5.380, control) vs (73.47 +/- 2.707 ChR2), n=8,8. **f,** Interaction time with familiar aggressor during 15 minute stimulation averaged across bins on day 10 of RSDS. Two-tailed Mann-Whitney T-test ***P=0.0002, (48.93 +/- 1.615, control) vs (64.67 +/- 1.905 ChR2), n=8,8. **g,** percentage change of time spent interacting with familiar aggressor between the 8^th^ and 10^th^ SDE at Bin 1 of day 8 compared to bin 8 of day 10. Two-way ANOVA (Bin x stimulation) interaction F(1,20) = 4.252 P=0.0524, time (Bins) F(1,20) = 4.038 P=0.0582, stimulation F(1,20) = 4.252 P=0.0524. **h,** Correlation of intra-defeat stimulation (familiar aggressor) by social interaction time on the SI test (novel non-aggressor conspecific). Control (n=8), linear regression goodness of fit test, control R^2^=0.2596, F (1,6) = 2.103, P=0.1972. **I**, Effects of intra-defeat stimulation on ChR2 and Control mice toward familiar aggressor over time, as a percent change in social interaction time from baseline (bin 1, no stimulation of day 8) to post-stimulation (bin 8, no stimulation of day 9). Two-Way ANOVA Interaction F(1,20) = 4.252 P=0.0524, Time F(1,20) = 4.038, P=0.0582, Control vs ChR2 F(1,20) = 4.252, P=0.0524. **j,** Correlation of intra-defeat stimulation (familiar aggressor) by social interaction time on the SI time. F(1,6) = 2.103, P=0.1972.

**Extended Data Fig 2:**
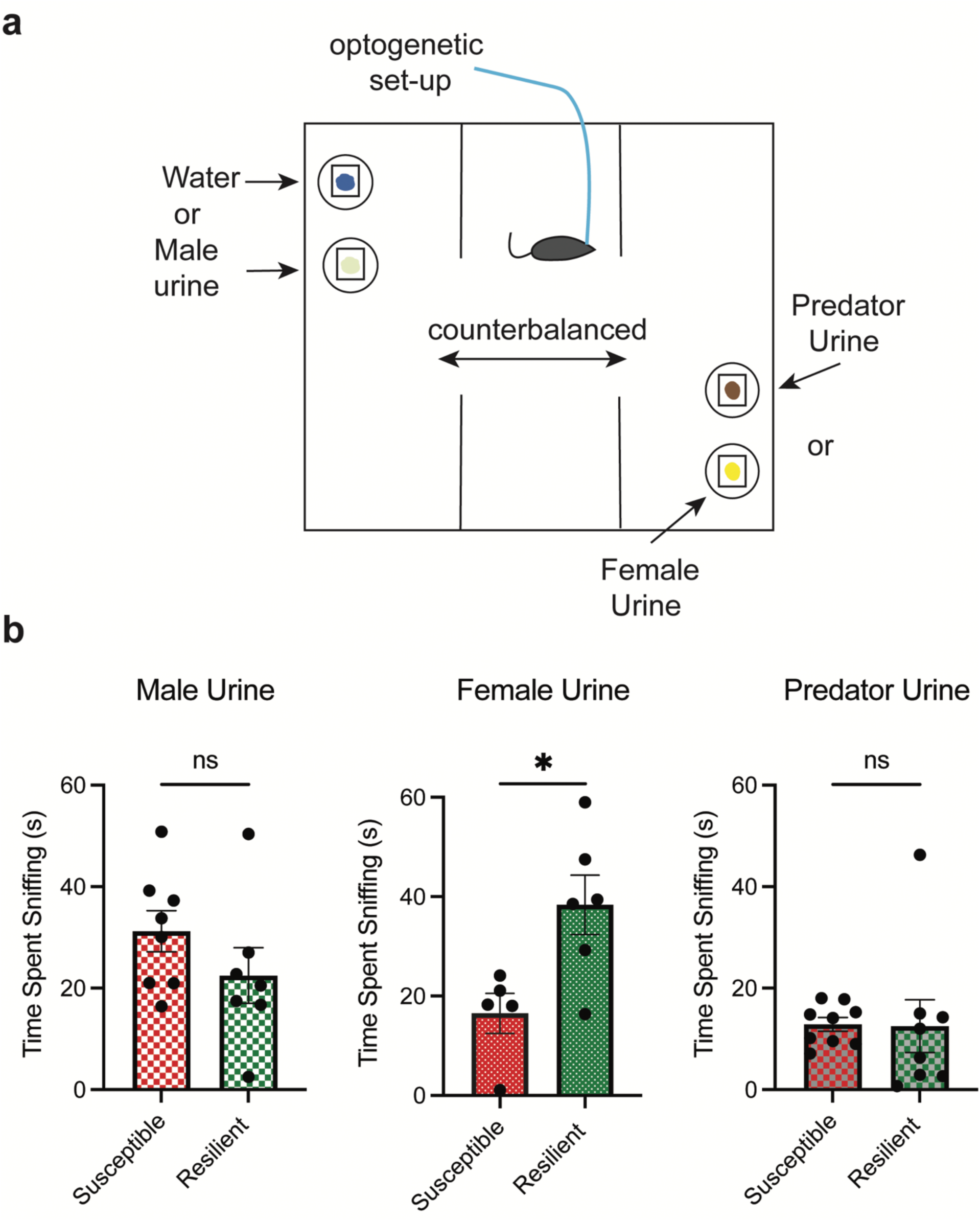
Distinct defeat-induced phenotypical differences convey individual preference for positively-valenced odors. **a,** Schematic of placement of odorants in three-chamber setup where mice are subjected to three trials contained odorants. **b,** Susceptible and resilient mice during “no light” trial interaction with male urine. T-test, two-tailed, P=0.2156, n=8 susceptible, n=7 resilient. **c,** Susceptible and resilient mice during “no light” trial interaction with female urine. T-test, two-tailed, *P=0.0178, n=5 susceptible, n=6 resilient. **d,** Susceptible and resilient mice during “no light” trial interaction with predator urine. T-test, two-tailed, *P=0.9454, n=9 susceptible, n=8 resilient.s

